# Codon biased translation mediated by Queuosine tRNA modification is essential for the virulence of *Leishmania mexicana*

**DOI:** 10.1101/2025.06.30.662252

**Authors:** Bankatesh Kumar, Julie Kovářová, Michala Boudová, Sneha Kulkarni, Thalia Pacheco Fernandez, Abhay Satoskar, Zdeněk Paris

## Abstract

The complex life cycle of the human parasite *Leishmania mexicana* requires rapid translational adaptation for survival in two distinct environments: the insect vector and the mammalian host. These protists lack conventional transcriptional control due to their unusual genome organization. Consequently, tRNA modifications may represent an additional mechanism for post-transcriptional regulation of gene expression. One such modification is queuosine (Q), which is incorporated at the anticodon wobble position 34 of specific tRNAs. Here, we demonstrate that Q-tRNA levels increase substantially during *Leishmania* differentiation from the insect stage to the mammalian-infective stage, implying an important role for virulence. Hence, we generated mutant cells lacking the enzyme responsible for Q incorporation, tRNA-guanine transglycosylase (TGT), which exhibited substantial changes in the proteome during differentiation *in vitro*. Specifically, downregulated proteins were enriched in NAU codons, whereas upregulated proteins predominantly contained NAC codons. Although LmxTGT knockout parasites exhibited normal growth and differentiation *in vitro*, they demonstrated impaired survival within macrophages and reduced pathogenicity in mice, highlighting the role of the Q-tRNAs under stress conditions. To our knowledge, we present here the first direct evidence that queuosine tRNA modification controls the infectivity of *Leishmania* via codon-biased translation. To date, gene expression regulation in *Leishmania* and other trypanosomatids, has been attributed mostly to RNA stability and processing, however, our findings demonstrate that tRNA modifications also play a key regulatory role. Specifically, the Q-tRNA modification provides a novel layer of gene expression regulation, maintaining translational balance and supporting the parasite’s ability to adapt to changing environments, and contributing to *Leishmania* virulence.

## INTRODUCTION

*Leishmania mexicana* (*L. mexicana*) is a kinetoplastid protozoan parasite and the primary cause of cutaneous leishmaniasis in North and Central America (1, 2). Leishmaniasis is a significant public health issue due to its high morbidity, causing skin lesions and long-term disability (1–3). Despite this, there are no approved vaccines for humans, and existing treatments often carry severe toxicity risks and can lead to parasite resistance (4–6). *Leishmania* parasites are transmitted to mammalian skin tissue during a blood meal via saliva of an infected sand fly. As a result, metacyclic promastigotes, which represent the infectious stage of *L. mexicana*, are rapidly phagocytosed by macrophages recruited to the site of the sand fly bite. Within the phagolysosomal compartment of these host cells, the parasites differentiate into the amastigote stage and proliferate intracellularly. During a subsequent blood meal, infected macrophages are taken up by another sand fly, continuing the transmission cycle (7). The progression of *Leishmania* life cycle involves a series of differentiation processes, including rapid remodelling of cellular architecture and physiological properties that enable adaptation to the environmental changes (8, 9). *Leishmania* and other trypanosomatids, such as *Trypanosoma brucei* and *Trypanosoma cruzi*, share distinctive mechanisms for regulation of gene expression (9). Unlike many eukaryotes, functionally unrelated genes in these parasites are organised into large, polycistronic transcription units that lack conventional promoter sequences. Transcription by RNA polymerase II begins in strand-switch regions, producing polycistronic pre-mRNAs (10). These are processed into mature mRNAs through trans-splicing, which appends a splice leader RNA at the 5’ end, and polyadenylation at the 3’ end. Consequently, gene expression regulation is largely post-transcriptional, involving mRNA stability, processing, and translation efficiency mediated for instance by 3’ UTR sequences (11–13), however, control of these processes is still not fully understood.

Another potential mechanism for fine-tuning of gene expression at the post-transcriptional level involves tRNA modifications that affect pairing at the wobble position (position 34). Depending on their chemical properties, these modifications can alter codon recognition potential and directly influence translational efficiency and fidelity, thereby affecting overall gene expression (14). One such modified nucleoside is queuosine (Q), a hypermodified analogue of guanosine (G), which is found in tRNAs with the GUN anticodon sequence, including those for asparagine (Asn), aspartic acid (Asp), histidine (His), and tyrosine (Tyr) (15). Q is present in all domains of life, but although bacteria can synthesise it *de novo*, this biosynthetic pathway is absent in eukaryotes. Therefore, they acquire the fully modified free base queuine (q) from their diet or the gut microbiome (16, 17) or salvage it in the cytosol through queuine-nucleoside hydrolase (QNG1), which mediates the recovery of queuine from queuosine-5′-monophosphate as the biological substrate (18). In eukaryotes, the formation of Q-tRNAs involves an isoenergetic reaction catalysed by the highly conserved heterodimeric enzyme tRNA-guanine transglycosylase (TGT) (19–21) (Fig. 1A), which breaks the glycosidic bond between the base and the sugar to replace the encoded guanine at position 34 with q.

**Figure 1:**
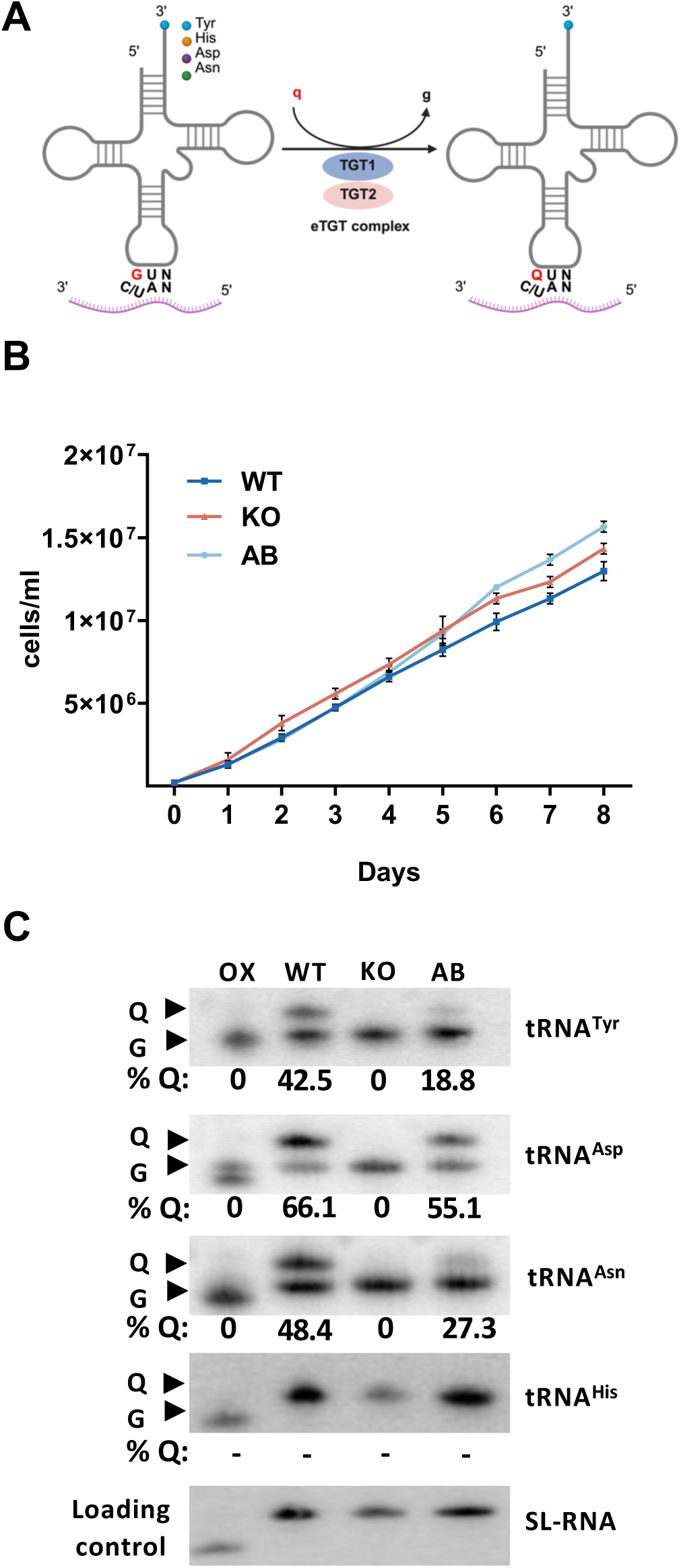
LmxTGT2 is necessary for Q-tRNA formation. **(A)** Schematic representation of the formation of queuosine (Q) tRNA modification in eukaryotes by the heterodimeric tRNA-guanine transglycosylase (TGT) complex, which catalyses the replacement of guanine (g) with queuine (q) in four specific tRNAs. (Created with BioRender.com) **(B)** Growth curves of wild-type (WT), LmxTGT2-KO (KO), and add-back (AB) promastigote cells exhibit no differences between the tested cell lines. (n = 3) **(C)** Northern hybridization showing the effect of LmxTGT2 gene knockout on Q-modified tRNA levels. Total RNA from WT, LmxTGT2-KO, and AB cells was separated on an APB affinity gel and hybridized with ³²P-labelled tRNA specific probes. As a result, Q-tRNAs migrate slower on the gel, leading to separation of modified (Q) and unmodified (G) tRNAs. Periodate oxidation (OX), which eliminates the mobility shift by oxidizing *cis*-diols (queuosine as well as terminal 3’ ribose), serves as a negative control. The splice leader (SL) RNA is used as a loading control. The numbers below the blot indicate the percentage of Q-modified tRNA, calculated by dividing the intensity of the Q band by the sum of the Q and G bands and multiplying by 100. The results are representative of three independent experiments.

Several studies point to the involvement of Q-tRNAs in processes such as cell proliferation, differentiation, or virulence (15, 22, 23). The position of Q at the wobble base of the anticodon of tRNA strongly implicates its role in translation via a mechanism based on codon usage. It has been suggested that Q-tRNA plays a crucial role in translation by modulating codon-anticodon interactions. *In silico* based analysis has shown that while unmodified tRNAs prefer NAC codons over NAU codons, Q-tRNAs accelerate translational speed at NAU codons (24). Further, *in vivo*, Q34 in cytoplasmic tRNAs was shown to regulate elongation speed, ensuring proper folding of nascent proteins and maintaining proteome integrity (17). In addition to its cytoplasmic role (25, 26), Q34 is also involved in mitochondrial translation, where it facilitates efficient NAU decoding in human mitochondria (27). These findings are in line with our previous observations in *T. brucei*, where Q-modified tRNAs are preferentially imported into the mitochondria, playing a crucial role in organellar protein synthesis (28). Without the Q modification, mitochondrial translation is impaired, leading to reduced activity of respiratory complexes and highlighting the importance of Q modification for organellar physiology (28). While the predominant trend seen in mammals (26), *T. brucei* (28), and bacteria (32), suggests that Q-tRNA preferentially improves decoding of Q-dependent codons ending with U over those ending with C, a few exceptions exist. In *Drosophila,* Q-tRNA demonstrate a bias toward NAC codons (30). Similarly, in yeast *Schizosaccharomyces pombe*, Q modification has been shown to regulate translation speed by enhancing the decoding of C-ending codons for aspartate and histidine while reducing the speed of U-ending codons for asparagine and tyrosine (31). Moreover, the position and sequence context of the impacted codons may be also important.

All these findings underscore the regulatory potential of Q-tRNA to modulate codon usage bias and translation efficiency, which could have significant implications for the adaptability and virulence of protozoan parasites such as *L. mexicana*. Additionally, recent research in mammal’s links Q-tRNA to translation fidelity, with its loss associated with impaired translation elongation and changes in neuronal morphology (32). Similarly, in human cells lacking queuine, codon usage imbalances have been observed in genes linked to differential protein expression, further emphasising the regulatory role of Q in translation (26, 31). While Q-tRNA modification has been associated with biofilm formation and virulence in bacteria (29), highlighting its role in pathogenesis, the impact on virulence in eukaryotic pathogens remains largely unexplored.

In this study, we investigated the role of queuosine tRNA modification in *L. mexicana* and its role in parasite differentiation and virulence. Using CRISPR/Cas9, we generated LmxTGT2 knockout parasites and established that Q-tRNA modification influences codon-biased translation, specifically affecting the expression of genes enriched in NAU codons. We further examined the functional consequences of Q-tRNA depletion in both macrophage infections and an *in vivo* mouse model. While LmxTGT2 knockout cells exhibited normal growth and differentiation when cultivated *in vitro*, their ability to survive within macrophages was compromised, and they demonstrated markedly decreased virulence in mice. Our findings highlight the crucial role of queuosine tRNA modification in regulating *Leishmania* translation and consequently its virulence to the mammalian host.

## MATERIALS AND METHODS

### Cell Culture

The three cell lines wild type (WT), LmxTGT2 knock-out (KO), and add-back (AB) strains of *L. mexicana* were grown in M199 medium (Sigma-Aldrich, Cat. No. M0393-10X1L), which was supplemented with 2 μg/ml biopterin (Sigma-Aldrich, Cat No. B2517-25MG), 2 μg/ml hemin (Sigma-Aldrich, Cat No. 51280-5G), 25 mM HEPES (Applichem, Cat. No. A3268), 100 units/ml penicillin, 100 μg/ml streptomycin (Sigma-Aldrich, Cat. No. P4333-100ML), 10 % fetal bovine serum (FBS) (Sigma Aldrich, Cat. No. F7524-500ML), and 100 μg/ml hygromycin (Invitrogen, Cat. No. ant-hg-1) at 25°C. The medium used for the cultivation of the LmxTGT2-KO cell line was also supplemented with 100 μg/ml nourseothricin (Jena Bioscience, Cat No. AB-102L) and 50 μg/ml puromycin (Invitrogen, Cat No. ant-pr-5) serving as selection markers. To determine the cell density, samples were fixed with a LeishFix solution (3.6 % formaldehyde, 150 mM NaCl, 15 mM Na3C6H5O7) and cell count was determined using a Neubauer hemocytometer (Sigma Aldrich, Cat No. BR717805-1EA). Bone marrow-derived macrophages were cultured in DMEM/F12 medium (Invitrogen, Cat No. 10565), supplemented with 100 units/ml penicillin and 100 μg/ml streptomycin (Sigma Aldrich, Cat. No. P4333-100ML) and 10 % FBS at 37°C (prior to infection) or 34°C (post infection) and 5 % CO2.

### Generation of LmxTGT2-KO and AB cell lines

The LmxTGT2-KO cell line was generated by CRISPR/Cas9 approach using the cloning strategy described by Ishemgulova et al (33). The LmxTGT2 gene specific single guide RNA (sgRNA) (sequence: GAAGATGGTGCAGGTAAGCG│CGG) was designed using the eukaryotic pathogen CRISPR guide RNA/DNA design tool (Peng & Tarleton, 2015; http://grna.ctegd.uga.edu), amplified by PCR using specific primers (Suppl. Table 1), and cloned into a modified pLEXSY-SAT2.1 vector (Jena Bioscience, Cat. No. EGE-274) containing a U6 promoter and a nourseothricin resistance marker. The vector was kindly provided by Prof. Vyacheslav Yurchenko (Life Science Research Centre, Faculty of Science, University of Ostrava). The repair template was PCR-amplified using corresponding oligonucleotides (Suppl. Table 1), generating a cassette containing the puromycin resistance gene flanked by 30 bp LmxTGT2 homology arms at the Cas9 cleavage sites. Ten micrograms of both the sgRNA construct and repair template were co-transfected into the *L. mexicana* Cas9-expressing cell line using the Amaxa Nucleofector™ program U-033. LmxTGT2 ablation was confirmed by PCR on genomic DNA using specific primers (Suppl. Table 1). The addback cell line was generated using expressing a Cas9-resistant version of the LmxTGT2 gene cloned in a modified pLEXSY vector, which contains UTRs of the A600 gene, a constitutively expressed throughout all parasite life stages.

### Northern blot analysis

Total RNA was isolated by phenol-chloroform extraction as previously described (34). For aminophenylboronic acid (APB) affinity electrophoresis, total RNA samples were deacylated by incubating them in 100 mM Tris (pH 9.0) for 30 minutes. To create an oxidation control, RNA was further treated with 2 mM NaIO₄ in 50 mM Tris (pH 5.0) in the dark for 2 hours at 37°C, preventing APB binding by modifying the ribose hydroxyl groups. The oxidation reaction was subsequently quenched with 2.5 mM glucose before use. A 10 ml gel mixture (8 M urea, 8 % polyacrylamide) was supplemented with 25 mg of APB (Sigma-Aldrich) before polymerisation (35). Electrophoresis of APB gels was conducted for approximately 6 hours at 75 V and 4°C, followed by electroblotting onto Zeta-Probe membranes under standard transfer conditions. The membrane was then UV-cross-linked for 1 min. For hybridisation, γ-32P ATP-labeled oligonucleotide probes were prepared using T4 Polynucleotide Kinase (NEB, Cat. No. M0201S) and purified using a Sephadex™ G-25 column. The probes were denatured at 98°C for 5 minutes before being added to the hybridisation buffer (5× SSC, 20 mM phosphate buffer, 7 % SDS, 1× Denhardt’s solution, and 1 mg/ml salmon sperm DNA). The membrane was hybridised at 48°C overnight, then washed twice at 48°C (Wash I: 3 × SSC, 5 % SDS, 25 mM NaH₂PO₄, 10× Denhardt’s solution; Wash II: 1× SSC, 1 % SDS). Hybridised membranes were exposed overnight to a phosphor-imager screen (GE Healthcare) and analysed using a Typhoon FLA 9000 scanner with ImageQuant TL software (GE Healthcare). Probes utilised for Northern hybridisation are listed in supplementary Table 1.

### *In vitro* differentiation

To induce differentiation of *L. mexicana* from the procyclic promastigote stage into infective metacyclic forms, cultures were initiated at a density of 2 × 10⁵ cells/ml and maintained in M199 medium (pH 7) supplemented with 10 % FBS for 10 days without dilution. Amastigote forms were obtained by transferring the metacyclic culture into SDM medium containing 20 % FBS and incubating at 32°C for an additional four days (36)

### qPCR

To assess stage-specific gene expression by quantitative reverse transcription PCR (qPCR), RNA was harvested on days 0, 8, 10, 13, and 17 of differentiation. Reverse transcription was performed using the QuantiTect Reverse Transcription Kit (Qiagen, Cat. No. 205311). cDNA was then used for qPCR with 60S primers as a control, PfrD1 primers, Sherp primers and Amastin primers (Suppl. Table 1). Reactions were carried out using Power SYBR™ Green PCR Master Mix (Thermo Fisher Scientific, Cat. No. 4367659) according to the manufacturer’s instructions and executed on the QuantStudio 6 PCR system (Thermo Fisher Scientific).

### Macrophage isolation and *in vitro* infection

Bone marrow-derived macrophages were isolated from BALB/c mice as described previously (37). Murine macrophages were infected with *L. mexicana* parasites following a previously described protocol, with minor modifications (37). Prior to infection, differentiated metacyclic *L. mexicana* were labelled with Cell Tracker^TM^ Orange CMRA dye (Invitrogen, Cat No. C34551) as described by Michael *et. al* (38). Macrophages were primed with IFN-γ (5 ng/ml; Invitrogen, Cat. No. PMC4031) 24 hours prior to infection and subsequently activated with a combination of IFN-γ (5 ng/ml) and LPS (10 ng/ml; Sigma-Aldrich, Cat. No. L2630-10MG). Subsequently, macrophage cultures were exposed to parasites at 1 to 7 ratio in a final volume of 2 ml for 4 hours at 34°C, 5 % CO2. The non-internalised parasites were removed with four successive washes with 1x PBS and subsequently replaced with fresh medium. This point was considered the initial time of infection (day 0). Cultures of infected macrophages were incubated for up to 5 days under the same conditions. To assess infection rates, cells were seeded onto coverslips placed in 6-well plates. At designated time points (days: 0, 1, 2, 3, and 5), coverslips were removed, and cells were fixed and permeabilized with ice-cold methanol for 10 minutes. Following fixation, cells were washed three times with 1x PBS to remove residual methanol. Subsequently, cells were stained with DAPI (Thermo Fisher Scientific, Cat No. P36941). Infection rates were determined by counting 100 cells per group under a fluorescence microscope, identifying nuclei stained with DAPI and parasites labeled with CMRA dye (Thermo Fisher Scientific, Cat No. C34551).

### Dual luciferase reporter constructs design and assay

The Renilla luciferase (Rluc) sequence remained unaltered as an internal control, while the cognate codons of Q-tRNAs in firefly luciferase (Fluc) were either left unchanged (50 % NAC, 50 % NAU) or modified to exclusively C-ending (100 % NAC) or U-ending (100 % NAU) codons. These two recoded constructs were synthesized by Eurofins Genomics. All three reporter constructs were cloned into the pLEXSY vector under the regulation of the A600 UTR and introduced into *L. mexicana* WT and LmxTGT2-KO cell lines via electroporation. Positive clones were selected using puromycin. The reporter assay was performed using Promega Dual Luciferase kit, as per manufacturer’s protocol. Briefly, 2 × 10⁴ cells for the WT background and 2 × 10^5^ cells for the Lmx-TGT2-KO background were washed with PBS and resuspended in Passive Lysis Buffer (PLB) in a 96-well transparent flat-bottom plate. The plate was incubated at room temperature for 15 minutes with continuous shaking. Firefly luciferase activity (FLuc) was measured following the addition of the LARII substrate using a Tecan Spark luminometer. Subsequently, the Stop & Glo reagent, which quenches Firefly luminescence while activating Renilla luminescence, was added, and RLuc activity was recorded. The FLuc/RLuc ratio was calculated, and values were normalised to the control construct (50 % NAC + 50 % NAU).

### Proteomics

WT, LmxTGT2-KO, and AB cells were collected at distinct differentiation time points (Days: 0, 8, 10, 13, and 17) and analysed at the CEITEC Proteomics Facility, Brno, Czech Republic. Briefly, proteins extracted in SDT buffer were processed using filter-aided sample preparation as described elsewhere (39). The resulting peptides were analysed by liquid chromatography–tandem mass spectrometry (LC-MS/MS) using the nanoElute system (Bruker) coupled to the timsTOF Pro 2 spectrometer (Bruker). Prior to LC separation, tryptic digests were online concentrated and desalted using a trapping column (Acclaim PepMap 100 C18, Thermo Fisher Scientific) and eluted onto an analytical column (Aurora C18, Ion Opticks) via a 90-minute linear gradient. MS/MS data were acquired in data-independent acquisition (DIA) mode with a precursor isolation range of m/z 400-1012. DIA data were processed using DIA-NN (version 1.8.1) (40) in library-free mode against the UniProtKB *L. mexicana* proteome database (UP000007259). Protein MaxLFQ intensities were further processed in a containerized computational environment (OmicsWorkflows, version 4.7.7a), following standard preprocessing, log2 transformation, and differential expression analysis using LIMMA. Mass spectrometry proteomics data have been deposited in the ProteomeXchange Consortium via PRIDE (41) partner repository with the dataset identifier PXD061534.

### Proteomic Data Analysis and GO Enrichment Methodology

Proteomic data analysis was performed using R for visualization, including Venn diagrams and Gene Ontology (GO) term plots. Differentially expressed proteins were identified by comparing LmxTGT2-KO with WT and AB across multiple time points, focusing on those consistently up- or down-regulated during differentiation. GO enrichment analysis, conducted using TriTrypDB (https://tritrypdb.org), assessed the functional impact of these proteins by mapping GO terms to relevant biological processes.

### Determination of codon bias in proteomics data

To evaluate the impact of codon bias on protein expression, proteins significantly up- or downregulated in the proteomics data were analysed for codon usage of tyrosine (Tyr), asparagine (Asn), aspartate (Asp), and histidine (His). The corresponding mRNA sequences were retrieved from the NCBI database using UniProt IDs. For each coding sequence (CDS), the number of Q-containing codons for each amino acid was calculated as the “observed codon frequency”. The Z-score for codon usage was determined using the formula: z-score of codon = (Observed frequency of codon - Expected frequency of codons)/ Standard deviation of observed frequency of codon, where the expected frequency represents the genome-wide average frequency of the respective codon. Genome codon frequencies were obtained from DNA Hive (42). Analysis was performed in RStudio (version 4.3.3), with ggplot2 used for data visualization.

### Ethical statement

Mice were housed and handled in the animal facility of the Biology Centre in accordance with institutional guidelines and Czech legislation (Act No. 246/1992 and 359/2012). All experiments were approved and conducted under permit No. PP 56_2022P from the Czech Ministry of Environment.

### Mice infection

Six-week-old BALB/c female mice were subcutaneously inoculated in the right ear with 1 × 10⁶ of WT, LmxTGT2-KO, and AB strains of metacyclic *L. mexicana*. M199 medium, serving as a blank control, was injected into the ears of the control group mice. Each experimental replicate included six mice per group. Ear thickness was measured using a Teclock standard-type thickness gauge, with a range of 0 to 20 mm and a scale interval of 0.01 mm. Lesion size was monitored weekly. To determine the lesion thickness, the thickness of the uninfected ear was subtracted from the thickness of the infected ear for each experimental animal.

### Antibody ELISA

To prepare the antigen, *L. mexicana* parasites were washed three times with 1× PBS and subjected to three freeze-thaw cycles, alternating between liquid nitrogen and a 37°C water bath. The protein concentration was determined using a BCA assay (Pierce™ BCA Protein Assay Kit, Thermo Fisher Scientific). Medium-binding 96-well plates were coated with 5 μg/ml of freeze-thawed *L. mexicana* antigen and incubated overnight at 4°C. The following day, plates were washed with PBS-Tween (0.05 % Tween-20 in 1× PBS, pH 7.4) and blocked with 5 % milk in PBS-Tween for 1 hour at 37°C. Blood samples were collected from infected BALB/c mice via submandibular vein bleeding every two weeks. Serum was separated by centrifugation and used for the antibody ELISA assay. After washing with PBS-Tween, serum samples were added in duplicate to the first well, followed by serial dilution across the plate. The plates were incubated for 2 hours at 37°C, followed by washing. The plates were then incubated with HRP-conjugated antibodies IgG1 (BD Biosciences, Cat#559626, clone X56) or IgG2a (BD Biosciences, Cat#553391, clone R19-15) at a final concentration of 0.2 μl in 1 ml of PBS-FBS (3:1) for 1 hour at 37°C, washed again, and incubated with TMB solution (Fisher Scientific) until colour change was observed. The reaction was stopped with 5 % phosphoric acid. Absorbance was measured at 450 nm using a Molecular Devices SpectraMax M3 microplate reader, and SoftMax Pro software was used to determine antibody titers.

### Parasitic burden

For *in vivo* studies, ears and draining lymph nodes were collected at the time of harvest (12 weeks post-infection for BALB/c mice). The tissues were homogenized using a cell strainer in 3 ml of Schneider’s Insect medium (Gibco, USA) supplemented with 20 % heat-inactivated FBS and 1 % penicillin/streptomycin. Each ear was separated into two sheets of dermis, washed in PBS containing 2 % penicillin/streptomycin, and processed into small pieces. The tissue was then incubated in HBSS (Sigma-Aldrich, Cat: H9394) supplemented with 2 mM EDTA, 2 % FBS, and 1 % penicillin/streptomycin at 37°C for 30 minutes, shaking at 250 rpm. After centrifugation, the cell pellet was incubated in DMEM + 2 mg/ml Collagenase A + 5 % FBS + 1 % penicillin/streptomycin at 37°C for 1 hour, shaking at 250 rpm. The enzymatic reaction was stopped by adding FBS, and the tissue was mashed through a 70 μm strainer, then washed with DMEM + 20 % FBS + 1% penicillin/streptomycin. The resulting lymph node or ear cell suspensions were serially diluted (1:20) in duplicates in two 96-well plates, ensuring that each sample was diluted across 24 wells. After 7 days of incubation at 26°C, the plates were examined using an inverted microscope at 40x magnification. The values reported in the graphs represent the highest logarithmic dilution showing viable parasites.

## RESULTS

### The LmxTGT2 subunit is essential for the formation of Q-tRNAs in *L. mexicana*

The enzyme tRNA guanine transglycosylase was previously shown to consist of two subunits: the catalytic TGT1 and the regulatory TGT2(43, 44). We identified the LmxTGT1 (LmxM.29.1800) and LmxTGT2 (LmxM.27.0030) genes in the genome of *L. mexicana,* which show 67 % and 42 % identities to their orthologs from *T. brucei*, respectively. To study Q-tRNA modifications in *L. mexicana*, we focused on the LmxTGT2 subunit, as our previous studies in *T. brucei* demonstrated that both TbTGT1 and TbTGT2 subunits are essential for Q-tRNA formation (28). We generated a gene knockout of the LmxTGT2 subunit in the promastigotes of *L. mexicana* using CRISPR/Cas9 (Supplementary Fig. 1A). The absence of the LmxTGT2 gene was confirmed by PCR analysis of genomic DNA (Supplementary Fig. 1B). Subsequently, an add-back cell line was constructed using a pLEXSY vector containing a recoded Cas9-resistant copy of the LmxTGT2 gene flanked by the 5′ and 3′ untranslated regions (UTRs) of the A600 gene (LmxM.33.3640), which was previously reported to maintain stable expression throughout *Leishmania* differentiation (45).

To assess the effect of LmxTGT2 depletion, we monitored growth of the cultured procyclic promastigote stage, which showed no significant differences in growth rates among the wild type (WT), LmxTGT2 knockout (KO), and LmxTGT2 add-back (AB) strains (Fig. 1B). Next, we investigated the involvement of LmxTGT2 in the formation of the Q modification of tRNAs. Total RNA from WT, KO, and AB cell lines was analysed using acrylamidophenyl boronic acid (APB)-affinity electrophoresis, followed by Northern blotting (35), which allows separation of the Q-modified tRNAs from their unmodified counterparts. The steady-state levels of Q-tRNA modifications in WT procyclic promastigotes varied among different tRNAs: Q-tRNA^Asp^, Q-tRNA^Asn^, and Q-tRNA^Tyr^ showed modification levels ranging from 42 % to 66 %, whereas Q-tRNA^His^ was close to the detection limit of the assay.

The complete absence of the bands corresponding to Q-modified tRNAs in the LmxTGT2 KO strain, along with a recovery of 44 – 83 % of WT levels (depending on the tRNA species) observed in the AB strain (Fig. 1C), illustrates the direct role of LmxTGT2 in Q-tRNA formation.

### Q-tRNA levels increase during the differentiation from the promastigote to the amastigote stage

To determine whether the levels of Q-tRNA modification change during the life cycle and whether its depletion affects the development of the parasite, we induced differentiation in the axenic culture. To simulate differentiation *in vitro*, promastigote cultures are left proliferating until they reach stationary phase and transform into metacyclic promastigotes. Subsequently, cells are transferred into Schneider’s insect medium (pH 5.5) and incubated at 32°C, which induces further differentiation into the amastigote stage (36) (Fig. 2A). However, the absence of Q-tRNA did not result in any noticeable changes in the growth or morphology of the LmxTGT2-KO parasites when growth was assessed daily up to day 13, within the same flask, and compared to the WT and AB strains (Fig. 2B).

**Figure 2:**
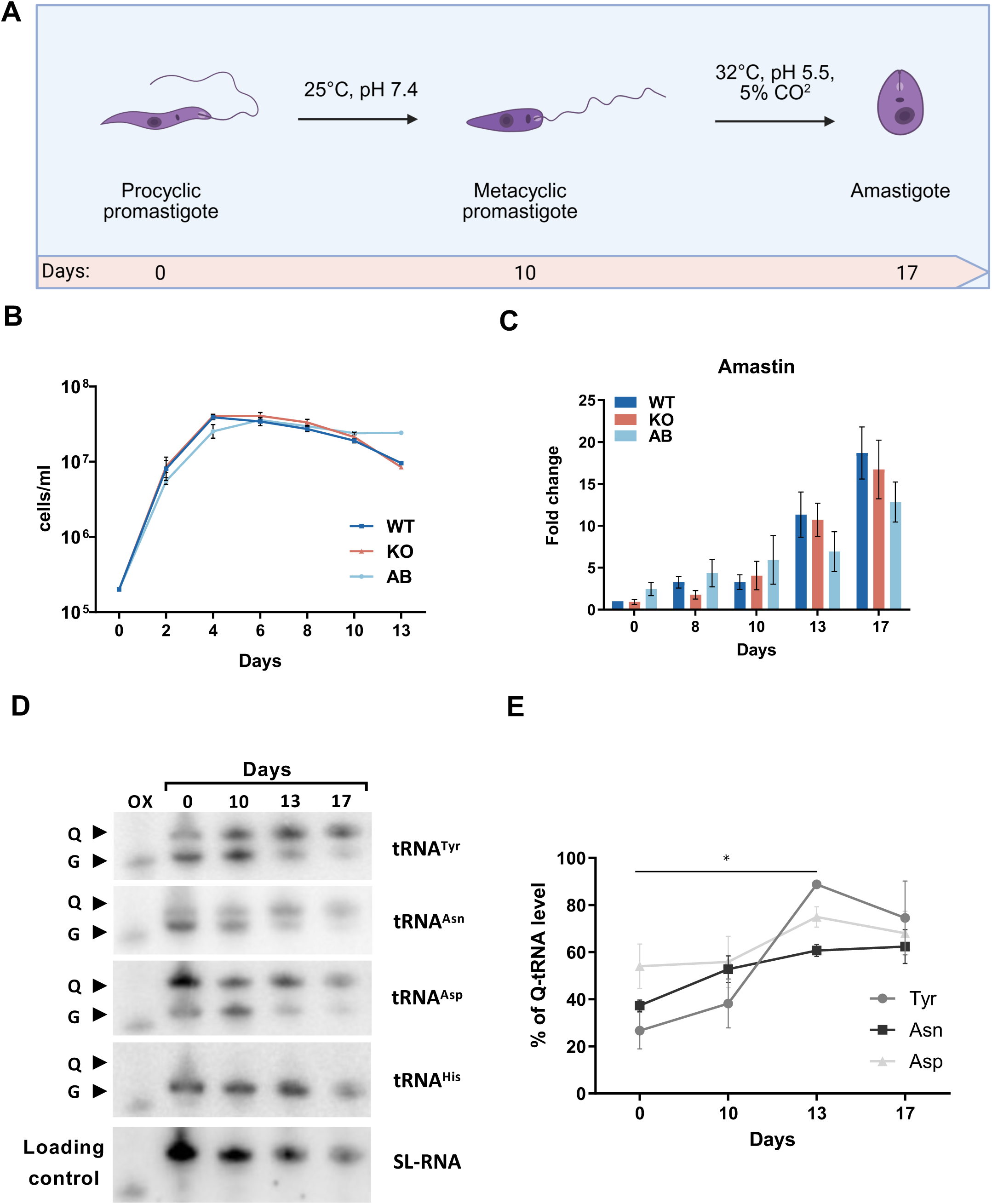
Q-tRNA levels increase during differentiation from the insect promastigote stage to the mammalian amastigote stage. **(A)** Schematic diagram of *in vitro* differentiation of *L. mexicana*. (Created with BioRender.com) **(B)** The growth curves of WT, LmxTGT2-KO, and AB cells exhibit no major alterations during *in vitro* differentiation into metacyclic promastigotes. Procyclic promastigotes were maintained in the same cultivation medium until reaching the stationary phase, at which point they differentiated into metacyclic promastigotes. (n = 3) **(C)** qPCR analysis of amastin marker expression during differentiation shows no significant differences between the tested cell lines, indicating unaltered differentiation into amastigotes. (n = 5) **(D)** Northern blot analysis of total RNA from cells collected at 0, 10, 13, and 17 days of differentiation, separated on an APB gel. The oxidized RNA was used as a negative control (OX), and the splice leader (SL) RNA was used as a loading control. **(E)** Quantification of Q-tRNA levels from the Northern blot experiments (as shown in D). Data were compared by two-way analysis of variance (ANOVA) with significant differences indicated (\**P* < 0.05, n = 3).

Although no obvious defects were observed during the differentiation in LmxTGT2-KO or AB, we validated the developmental progress by detecting stage-specific markers. As expected, expression of the amastigote-specific marker amastin increased during the differentiation into amastigotes. No significant changes in amastin expression were observed across the different cell lines (Fig. 2C). Similarly, no cell line-specific differences were observed in the expression of the other stage specific markers (Suppl. Fig. 2 and Suppl. Fig.3). Collectively, these findings imply that although Q-tRNA modification increases during the life cycle, the absence of LmxTGT2 and subsequent lack of this modification does not substantially impact the differentiation process *in vitro*.

**Figure 3:**
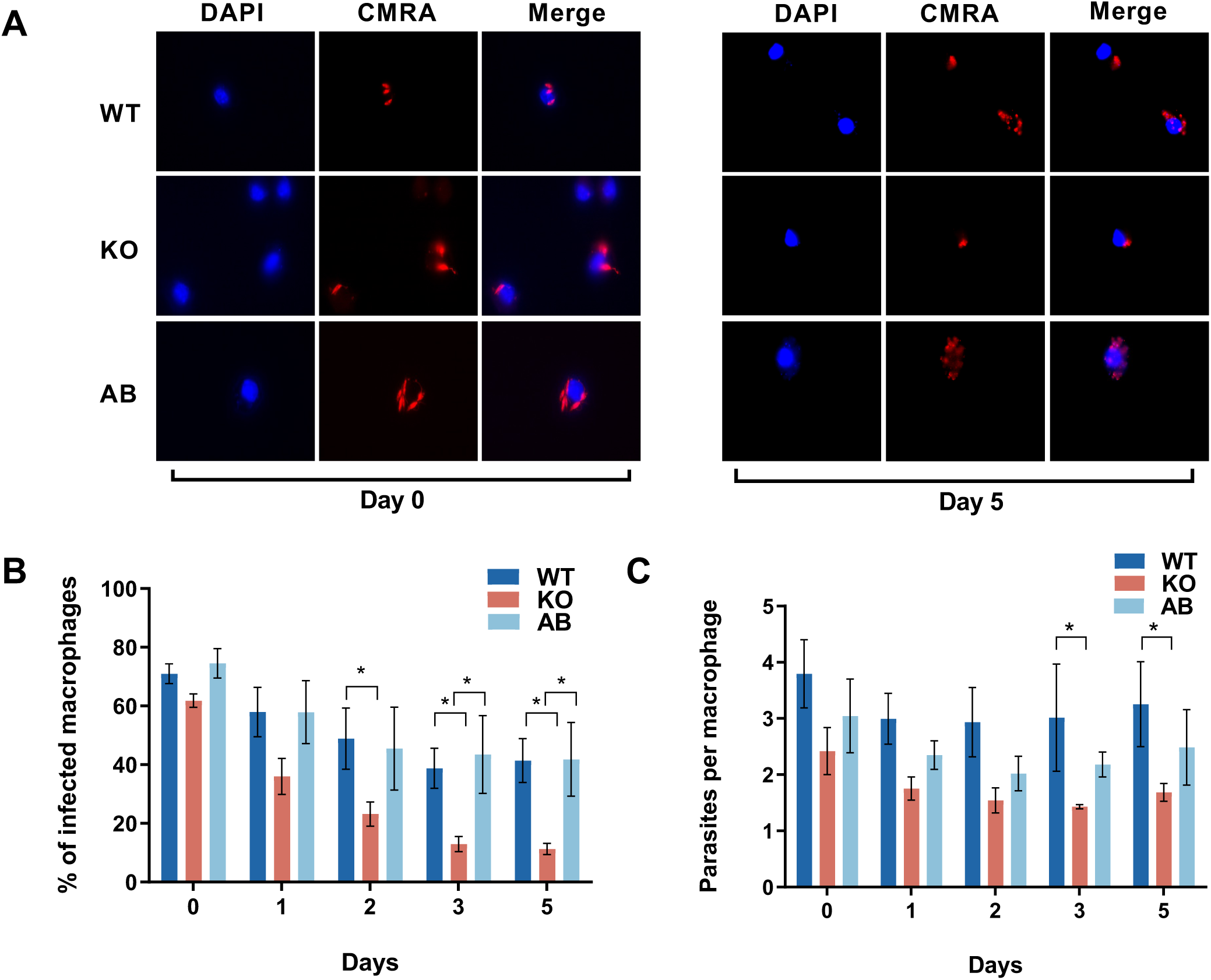
*L. mexicana* TGT2-KO has reduced infectivity and survival inside macrophages. **(A)** Fluorescence microscopy images of macrophages infected with WT, LmxTGT2-KO, and AB cells of *L. mexicana* are shown for illustration at day 0 and day 5 post-infection. Macrophage nuclei were stained with DAPI (blue), and parasites were labelled with CMRA orange cell tracker (red). **(B)** Quantification of macrophages infected with *L. mexicana*. Data represent three independent replicates of experimental infection assessed by fluorescence microscopy, showing the percentage of infected macrophages at days 0, 1, 2, 3, and 5 post-infections. A total of 100 macrophages were counted per sample. **(C)** The number of parasites per infected macrophage is shown. Data were compared by two-way analysis of variance (ANOVA) with significant differences indicated (\**P* < 0.05).

During the differentiation, we examined variations in the Q-tRNA modification steady-state levels across different life stages of the WT. Using APB electrophoresis of the total RNA, we assessed the Q content of tRNA^Tyr^, tRNA^Asn^, tRNA^Asp^, and tRNA^His^ at various differentiation time points: day 0, day 10 (metacyclic promastigotes), day 13 (early amastigotes), and day 17 (amastigotes). The results revealed a global increase in the Q-tRNA modification levels throughout the life cycle, peaking at the amastigote stage. The increase was the most prominent in the tRNA^Tyr^, and less striking but still prominent in tRNA^Asp^ and tRNA^Asn^ (Fig. 2D, E). Similar to the previous result (Fig. 1), Q modification in tRNA^His^ remained almost undetectable.

### LmxTGT2-KO compromises the ability of *L. mexicana* to infect macrophages *in vitro*

Although the lack of Q-tRNAs did not affect cell proliferation in the axenic cell culture, the levels of Q-tRNAs changed during differentiation, suggesting a possible importance in the infectious stages of the parasite. Thus, we hypothesised that the absence of Q-tRNAs could be more relevant if the parasites were subjected to a challenging environment within their mammalian host cells. To test the potential impact of LmxTGT2-KO on the infectivity of the parasites, we conducted an *in vitro* infection of bone marrow-derived murine macrophages. The differentiated parasites were fluorescently labelled, and infection was monitored over five days following the initial infection. At the start of the experiment, 60 – 70 % of macrophages were infected (day 0) across all groups. However, the difference between the three tested groups became significant from day 2 to day 5 of infection (Fig. 3A, B). Over time, the number of macrophages infected with LmxTGT2-KO parasites decreased substantially, resulting in only 11 % infected macrophages on day 5, compared to 41 % in the WT group (*P* = 0.0157) (Fig. 3B). Correspondingly, the number of parasites detected inside a macrophage was comparable at the beginning of the infection, but progressively decreased for LmxTGT2-KO parasites, while remaining unchanged for WT and AB cell lines (Fig. 3C). Overall, these data indicate that LmxTGT2-KO parasites can initially infect macrophages at the same level as WT, but they are unable to survive inside the host cells for several days.

### Q-tRNA is important for cytosolic codon-biased translation

To investigate the role of Q in cytosolic codon-biased translation, we employed the dual luciferase reporter system, where the *Renilla* luciferase (RLuc) is fused to firefly luciferase (FLuc) with a linker (Fig. 4A) (46). Since these enzymes use different substrates, it is possible to quantify differential expression of these enzymes proteins simultaneously, in a luminescence assay. We generated codon re-engineered constructs, where the RLuc sequence was kept unaltered, but in the FLuc sequence, the cognate codons for all four Q-containing tRNAs (tRNA^His^GUG, tRNA^Asp^GUC, tRNA^Asn^GUU, tRNA^Tyr^GUA) were changed either to all NAC (100 % NAC) or all NAU (100 % NAU) codons, as opposed to the control 50 % NAC + 50 % NAU FLuc, which contained approximately equal number of NAC and NAU codons. The total numbers of NAC and NAU codons are enumerated in Fig. 4B. We hypothesized based on available literature and our previous results in *T. brucei*, that Q might be more efficient than G, in decoding of U-ending codons (28). Thus, in the absence of Q-tRNAs we might observe a decreased translation of NAU luciferase, and subsequently, a decrease in the luminescence. We transfected these three constructs into WT and LmxTGT2-KO cells and measured Fluc luminescence, normalizing it to Rluc. The absence of LmxTGT2 resulted in an overall defect in translation, since luminescence dropped to 75 % (*P* = 0.01) in KO cells with the control construct (50 % NAC + 50 % NAU). While the NAC construct did not show significant differences in signal between WT and LmxTGT2-KO cells, the NAU construct exhibited a significant decrease in Fluc luminescence in the absence of LmxTGT2 (Fig. 4C). This suggests that NAU codons may be translated less efficiently without Q-tRNAs, as hypothesized above.

**Figure 4:**
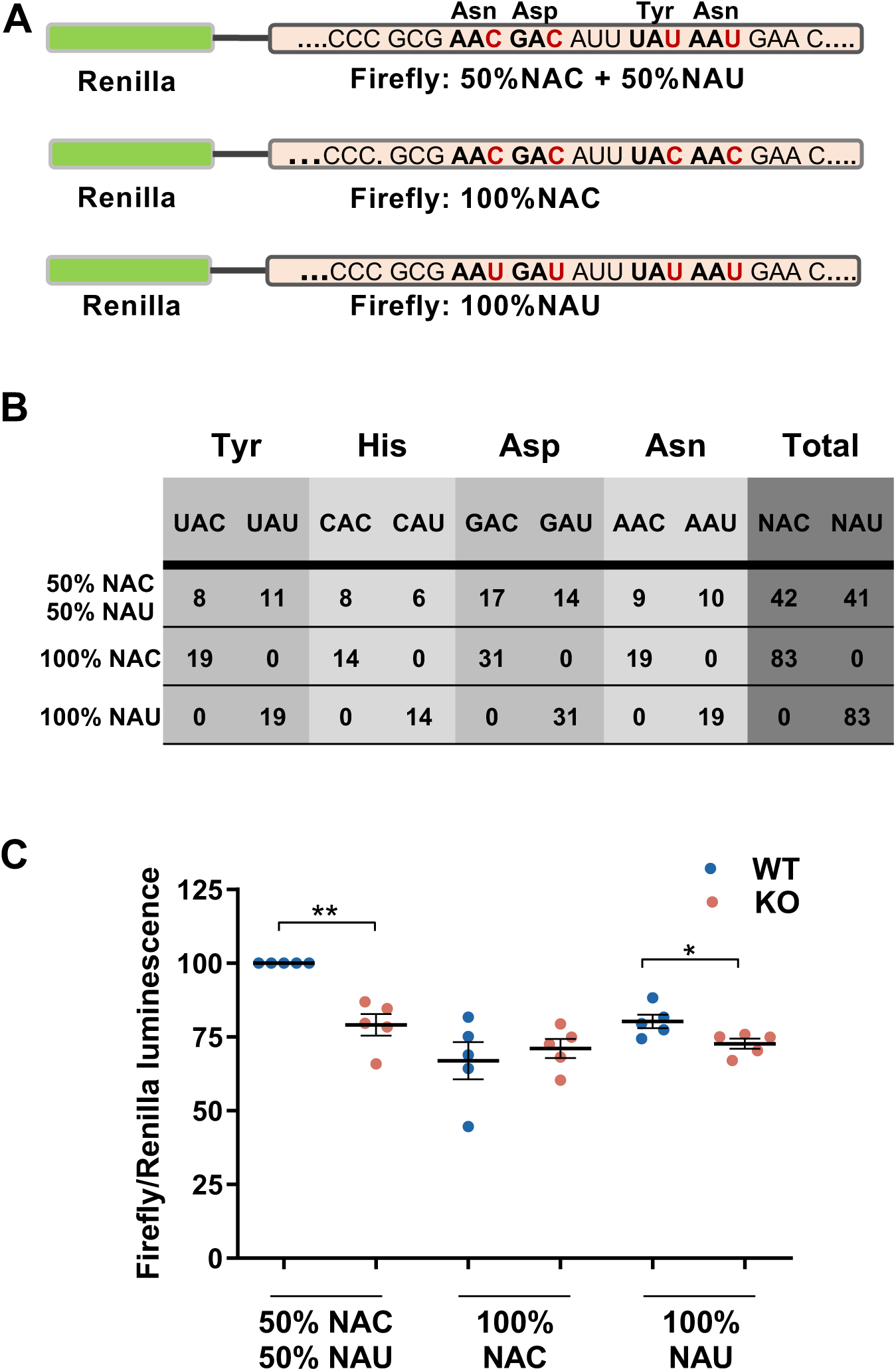
Q-tRNA is important for cytosolic codon-biased translation. **(A)** Schematic representation of a dual luciferase construct where *Renilla* luciferase (RLuc) is expressed in-frame with a codon-reengineered Firefly luciferase (FLuc). In the control construct, Q-dependent codons in the Fluc sequence are evenly distributed (50% NAC, 50% NAU), whereas in the test constructs, all Asn, Asp, Tyr, and His codons were recoded to contain exclusively either NAC or NAU (as shown in panel A). **(B)** Number of cognate codons for Q-tRNAs in the three test constructs. **(C)** Ratio of FLuc to RLuc luminescence measured in the cells, normalised to the control (50 % NAC + 50 % NAU), representing efficiency of translation of the reporter protein. Data were analysed using Student’s *t*-test, with significant differences indicated (\*\**P* < 0.01, \**P* < 0.05, n=5).

### The loss of Q-tRNA modification alters proteome of *L. mexicana* during differentiation

Since the luciferase assay indicated an effect of Q-tRNAs on codon biased translation, we performed a proteomic analysis to investigate the consequences of Q-tRNA depletion on the proteome during the *L. mexicana* life cycle. This analysis had two main objectives: first, to identify differentially expressed proteins in LmxTGT2-KO relative to WT and AB at different time points; and second, to examine proteins that showed consistent changes throughout differentiation in LmxTGT2-KO (relative to day 0). Our results revealed significant differences in proteome composition, particularly on days 8, 13, and 17 of the differentiation process (Fig. 5).

**Figure 5:**
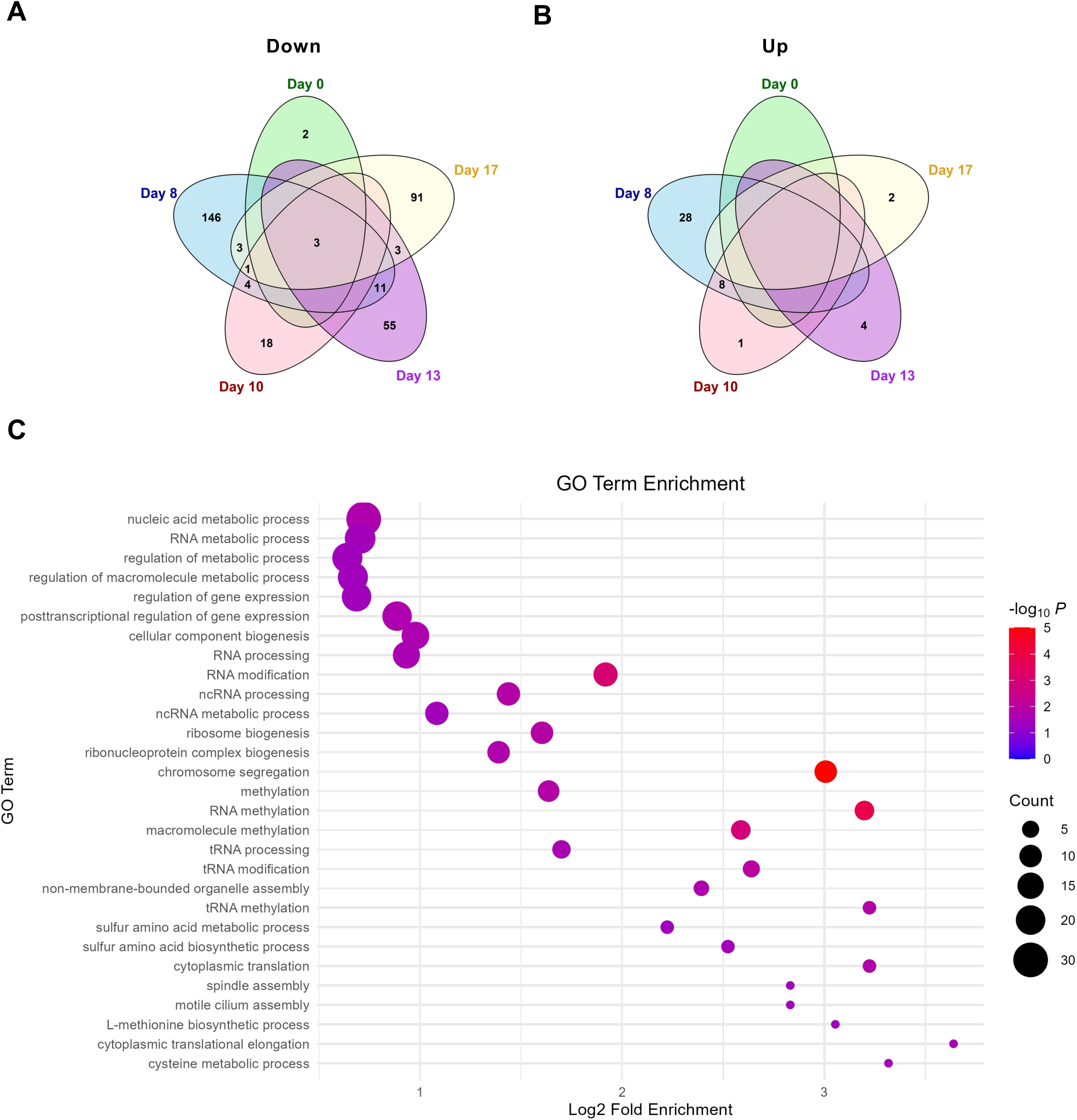
The absence of LmxTGT2 altered the protein profile during differentiation. **(A,B)** Venn diagrams showing the number of downregulated **(A)** and upregulated **(B)** proteins at different time points of differentiation in LmxTGT2*-*KO relative to WT and AB. In the absence of a number in a particular field, no common proteins were identified among the groups. Differential expression was defined as a fold change of ≥ 2 with statistical significance, as determined by Student’s *t*-test (*P* < 0.05, n=3). **(C)** Dot plot illustrating gene ontology (GO) enrichment analysis of proteins downregulated in LmxTGT2-KO cells throughout differentiation period (day 0 to day 17). GO terms with less than two associated proteins were excluded, resulting in the removal of 22 terms. Of the 344 downregulated proteins, 313 were assigned to GO terms, while 31 remained uncharacterized. The size of the dots indicates the number of proteins linked to each GO term, and the colour of the dots reflects the statistical significance of the enrichment.

In terms of the overall proteome, we identified 7236 proteins. In KO relative to WT and AB, we observed 168 proteins downregulated at day 8, 72 proteins at day 13, and 101 proteins at day 17. In contrast, fewer proteins were upregulated, with a peak of 36 upregulated proteins at day 8, followed by 9 proteins at day 10, 4 proteins at day 13, and just 2 proteins at day 17 (2-fold change, *P =* 0.05; Fig. 5A and 5B). Notably, there were not many common proteins among different time points. Interestingly, only three proteins LmxTGT1, LmxTGT2, and an uncharacterized protein, were consistently downregulated across all time points, forming the only overlap between all groups (Fig. 5A and 5B).

To assess proteomic alterations within LmxTGT2-KO during differentiation, we analysed proteins that were consistently downregulated or upregulated relative to the LmxTGT2-KO starting point (day 0), while remaining unchanged in WT and AB. Between day 8 and day 17 of differentiation, a total of 344 proteins were significantly downregulated, while 131 proteins were significantly upregulated (2-fold change, *P =* 0.05). To investigate the potential function of these downregulated proteins in LmxTGT2-KO cells, we performed the Gene Ontology (GO) enrichment analysis. Overall, 313 proteins were mapped to GO terms, while 22 GO terms were excluded due to having only one associated protein. The most significantly enriched GO terms were related to RNA metabolism and processing, including RNA metabolic processes, RNA modification, ribosome biogenesis, and non-coding RNA processing, suggesting a disruption in RNA processing. Additionally, processes related to gene expression regulation, posttranscriptional control, and methylation were notably affected. (Fig. 5C).

To further investigate the impact of Q-tRNA on translation, we examined the correlation between changes in protein expression and the usage of Q-specific codons, focusing on NAC and NAU codons. We observed a slight positive correlation between NAC codon frequency and protein levels on both day 8 (r = 0.166, *P* = 2×10^-16^) and day 13 (r = 0.149, *P* = 2×10^-16^). In contrast, NAU codons showed a trend towards negative correlation on day 8 (r² = -0.005, *P* = 0.653) and day 13 (r = -0.043, *P* = 0.000228) (Fig. 6A, Supplementary Fig. 4E).

**Figure 6:**
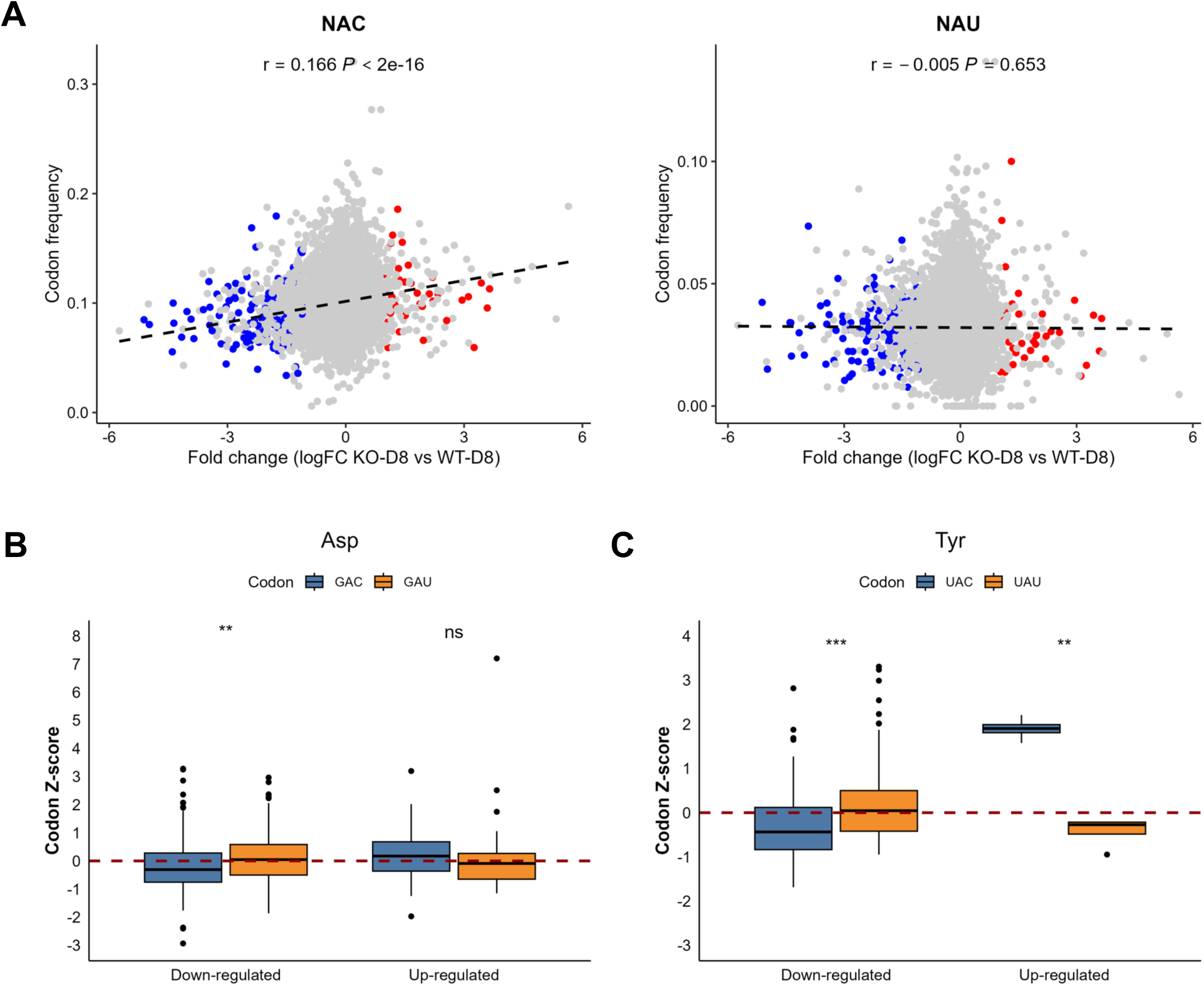
Q-tRNAs modulate translation during differentiation via codon bias. **(A)** Pearson correlation between codon frequency and protein fold change in upregulated and downregulated proteins for NAC and NAU codons in LmxTGT2-KO cells at day 8, relative to WT and AB. The Pearson correlation coefficient (r) with *P* value and linear regression line (black) are indicated. **(B)** Frequency of Asp codons at day 8 and Tyr codons at day 13 **(C)** was evaluated by calculating Z-scores for upregulated and downregulated proteins in KO cells. Here, the Z-score represents the degree to which observed codon usage deviates from the expected codon frequency in the genome. The center line indicates the median, the box shows the interquartile range (upper and lower quartiles), and individual dots represent outliers (minimum and maximum values). The data are based on three independent biological replicates used for proteomic analysis and were analysed using Student’s *t*-test. Significant differences are indicated (\*\*\**P* < 0.001, \*\**P* < 0.01, * *P* < 0.05, ns = not significant).

Since the effect of Q-modification may be specific to individual tRNAs and could be obscured in the broader analysis of NAC and NAU codons, we next focused on the specific codons involved. This analysis concentrated on amino acids decoded by Q-tRNAs, specifically tyrosine (Tyr), asparagine (Asn), aspartate (Asp), and histidine (His). Codon usage deviations were evaluated using the Z-score (47), which compares the observed frequency of specific codons with their expected frequency based on genome wide codon usage. The analysis revealed significant differences in the usage of Asp codons on the proteome of day 8, since GAU codon is enriched in down-regulated proteins (Fig. 6B). Further, Tyr codons showed pronounced enrichment of UAU in down-regulated proteins, and enrichment of UAC in up-regulated proteins on day 13 (Fig. 6C). Differences in codon usage at other time points were less pronounced. In general, Q-tRNA-dependent U-ending codons were enriched in down-regulated proteins, whereas C-ending codons were more prevalent in up-regulated proteins, in line with the original hypothesis (Suppl. Figure 4 and 5).

### The ablation of LmxTGT2 impacts parasite infectivity and virulence in the mammalian host

To determine whether the decreased infection observed in macrophages could be replicated *in vivo* and prompted by the profound effects on the LmxTGT2-KO proteome, an infection study was performed in Balb/c mice to investigate a role of LmxTGT2 in *L. mexicana* virulence. Parasites were administered intradermally into an ear of each mouse. Lesions became apparent after 8 weeks in all groups and continued to increase in size. From the ninth week onward, notable variations in lesion size were observed, with mice infected with LmxTGT2-KO parasites developing lesions approximately half the size of those caused by WT and AB strains. This difference became statistically significant from week 10 onward (Fig. 7A).

**Figure 7.**
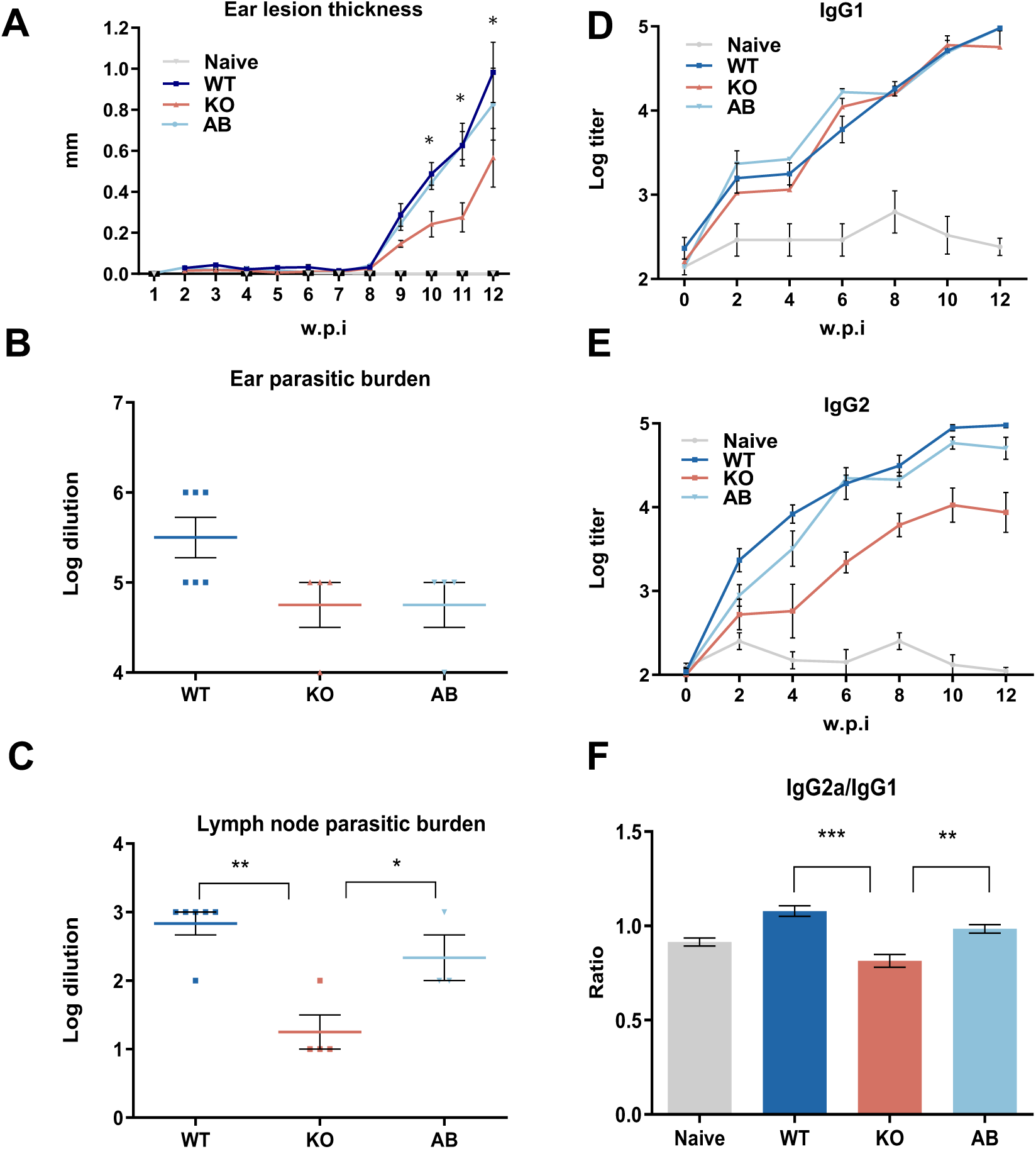
The ablation of LmxTGT2 impacts parasite infectivity and virulence in a mammalian host. **(A)** Balb/c mice were infected with *L. mexicana* wild-type (WT), add-back (AB), or LmxTGT2-KO parasites. Ear lesion size (in mm) was measured weekly over the course of infection (w.p.i. = weeks post-infection). Results shown are from one representative experiment out of two independent experiments. **(B)** Parasite burden in ear tissue, assessed by serial dilution of lesion-derived material at week 12 following the end of the infection experiment. **(C)** Parasite burden in the corresponding ear-draining lymph nodes (determined as in B). **(E)** Time course of the antibody response, showing IgG2a and **(D)** IgG1 levels measured by ELISA in sera from infected mice collected at two-week intervals. **(F)** IgG2a/IgG1 ratio was calculated from the data shown in panels D and E. Data were analyzed using Student’s *t*-test, and significance is indicated (** *P* < 0.01, \**P* < 0.05).

The assessment of parasite load through a serial dilution assay did not reveal significant changes in the number of parasites in the ear, despite the larger lesions observed in the WT and AB groups compared to LmxTGT2-KO (Fig. 7B). Interestingly, a lower number of parasites was detected in the lymph nodes in LmxTGT2-KO compared to WT and AB (Fig. 7C). To evaluate the inflammatory response in infected mice, we performed an enzyme-linked immunosorbent assay (ELISA) for IgG1 and IgG2a in the serum. Throughout the infection period, IgG1 levels steadily increased in all tested groups (Fig. 7D). In contrast, IgG2a titers were significantly lower in LmxTGT2-KO animals compared to WT and AB mice (Fig. 7E). The ratio of IgG2a to IgG1 was calculated to be significantly reduced in mice infected with the LmxTGT2-KO strain. This ratio is an important indicator of the immune response profile, where a higher IgG2a/IgG1 ratio typically reflects a Th1-biased (protective) immune response, whereas a lower ratio indicates a Th2 (anti-inflammatory) response (48, 49) (Fig. 7F).

## DISCUSSION

The wide range of biological phenomena associated with Q has made it challenging to pinpoint a single, unified role for this modification across species (17). Notably, intracellular transport dynamics and/or nutrient availability influence queuosine levels in *T. brucei* (25, 28, 50, 51). In addition, changes in Q-tRNA levels were observed in the insect and mammalian life stages of this trypanosomatid parasite (25). This conserved pattern of dynamic Q-tRNA modification has been observed in many other organisms, suggesting a broader regulatory role across eukaryotes and linking environmental cues and developmental programs to the fine-tuning of translation.

In this work, we show that in the protozoan parasite *L. mexicana*, Q-tRNA levels gradually increased during diÖerentiation, reaching their highest levels in the amastigote stage. This finding suggests that Q-tRNAs play a role in controlling stage-specific development. Indeed, deletion of the active subunit of TGT in *L. mexicana* (LmxTGT2) abolished Q-tRNA formation, but showed no apparent growth defects, suggesting that Q-modification may not be essential for basal proliferation and differentiation *in vitro,* but may be required under changing environment or nutrient-limiting conditions. This interpretation is consistent with findings in other systems, where deletion of TGT subunits (QTRT1 or QTRT2) did not impair proliferation under normal conditions (27, 52).

Q-tRNAs correspond to single tRNA isoacceptors decoding Asn, Asp, Tyr or His codons; all found in two-codon boxes (NAU and NAC). The N7-deaza guanosine-derived analog Q replaces guanosine at position 34 (wobble base), subtly altering base-pairing dynamics. While Q in most organisms does not alter the Watson–Crick face of the anticodon, it enhances wobble pairing with U over C, favoring decoding of U-ending codons in most contexts (17). However, in *S. pombe*, the opposite pattern has been reported, where Q preferentially enhances the translation of C-ending codons (31). As shown here, we observed a significant reduction in NAU-driven firefly luciferase activity in Q-tRNA deficient *L. mexicana* cells, analogous to our previous observations in *T. brucei* (28). However, even the NAC-only construct exhibited reduced translation regardless of the presence or absence of Q-tRNAs, suggesting that codon optimality itself contributes to translational output independent of Q-modification. Interestingly, constructs with a balanced 50:50 ratio of NAU and NAC codons also exhibited a reduction of translation in LmxTGT2-KO cells, indicating a general defect on translation in the absence of Q-tRNAs. A similar observation was reported in human cells lacking QTRT1 or QTRT2 (52).

Along these lines, our proteomic analysis revealed that proteins downregulated in Q-tRNA deficient cells were significantly enriched for NAU codons, whereas proteins upregulated under these conditions showed enrichment for NAC codons. Although codon usage alone does not fully explain the observed protein expression levels, these trends are consistent with the idea that Q-tRNA modification promotes efficient translation of NAU-rich transcripts. The trend becomes more pronounced when individual codon pairs are considered, suggesting that the effect of Q-modification varies depending on the specific tRNA involved.

As defined by Grosjean & Westhof (53), codons with A and/or U at the first two positions are thermodynamically “weak” whereas those with G and/or C are “strong”. In this context, Asn (AAU/AAC) and Tyr (UAU/UAC) codons are weak; Asp (GAU/GAC) and His (CAU/CAC) are intermediate. It has been proposed that Q-modified tRNAs may help buffer such stability differences, enhancing decoding kinetics across codon families with varying strength (53). While our study does not directly address this mechanism, the preferential reduction in translation of NAU-rich transcripts observed in Q-tRNA deficient cells is consistent with the idea that Q-tRNAs enhance decoding of weaker codons, fine-tuning decoding efficiency, which is particularly important for codons that are thermodynamically less favorable. Together, these data support a model where Q-tRNA modification facilitates codon-specific translation during developmental transition between the insect and mammalian stage (Fig. 8). Its absence compromises the efficient decoding of specific codons under changing environment, while remaining dispensable during homeostatic growth. Future work combining ribosome profiling with enhanced protein annotation will be required to resolve how Q-tRNA influences translation in a codon context and transcript-specific manner.

**Figure 8:**
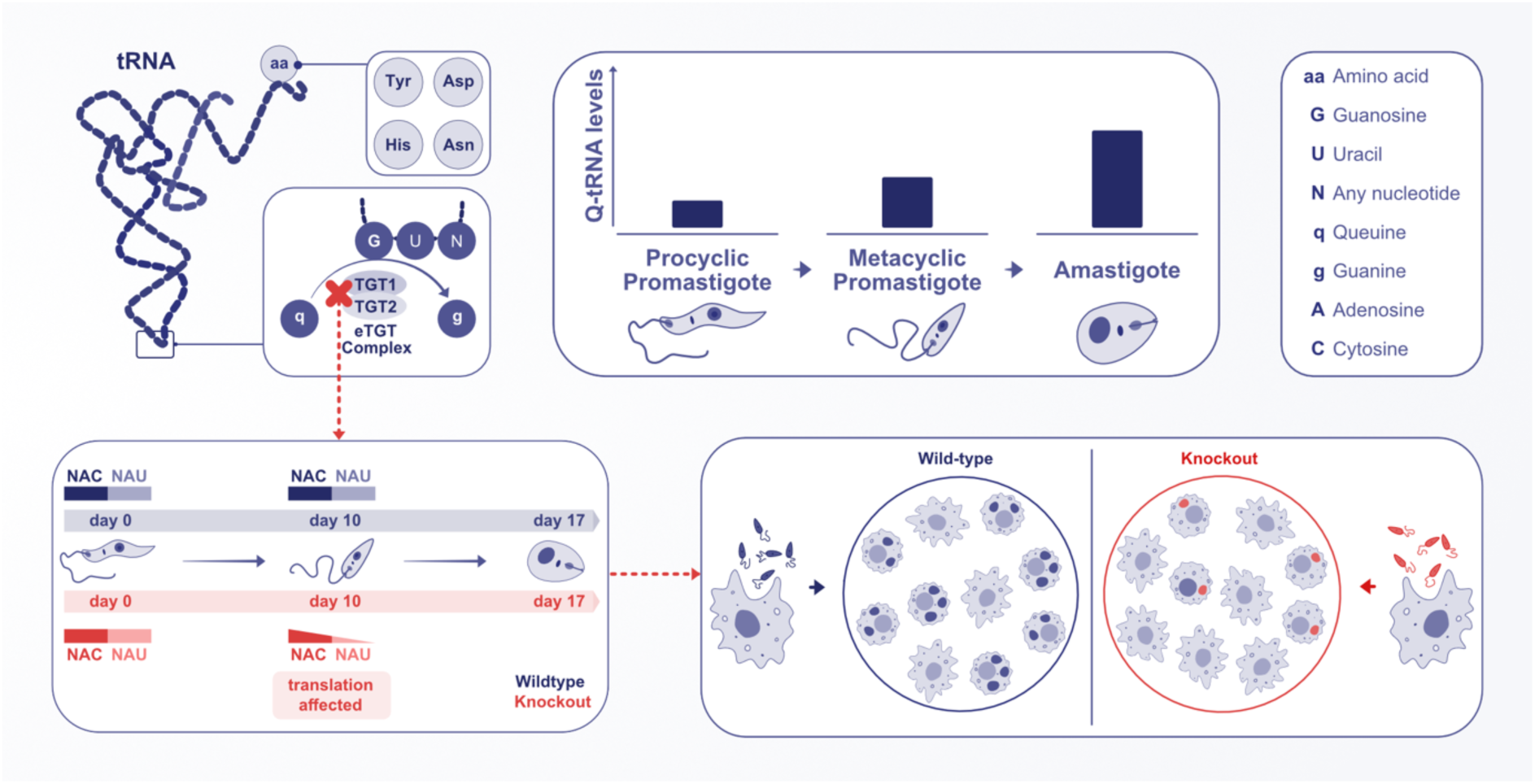
Model illustrating the role of queuosine (Q) tRNA modification in the virulence of *L. mexicana*. Q-tRNA levels increase significantly during the differentiation from the procyclic promastigotes (insect stage) to the amastigotes (mammalian stage), underscoring the importance of Q-tRNA modification for parasite adaptation and infectivity. Loss of Q modification via LmxTGT2 knockout disrupts codon-biased translational control, leading to substantial changes in the proteome. Specifically, downregulated proteins were enriched in NAU codons decoded by Q-tRNA, while upregulated proteins predominantly contained NAC codons. The resulting codon-specific imbalance impairs the parasite’s ability to establish infection and maintain virulence. These findings demonstrate that Q modification functions as a novel post-transcriptional regulatory mechanism that fine-tunes translation during *Leishmania* life cycle, compensating for the lack of classical transcriptional control and promoting adaptation to the host environment.

An open question is how Q-tRNA modification levels increase during *Leishmania* differentiation, given that under normal conditions tRNAs are not fully modified. One possible explanation is that the TGT complex itself becomes upregulated during differentiation; however, our proteomic data does not support this. Considering our previous observations in the *T. brucei* system (51), more likely scenarios may be either increased dwell time of tRNAs in the nucleus or changes in nutrient levels when cells develop into metacyclics and then into amastigotes, which are characterized by slower growth and reduced metabolic rates and higher sensitivity to perturbations (54). Notably, in the LmxTGT2 knockout, we observed a partial reduction in the overall levels of a subpopulation of tRNAs that typically undergo Q modification, consistent with previous findings that Q contributes to tRNA stability (55). This supports the interpretation that Q-tRNA depletion primarily exerts a subtle, codon-specific effect on translation, with broader translational impairments arising as a secondary consequence.

Several studies suggest that Q plays a role in modulating virulence across both prokaryotic and eukaryotic pathogens (29, 56–58). To our knowledge, the only eukaryotic pathogen that has been investigated in the context of Q is *E. histolytica*, where queuine supplementation slightly increased Q-tRNA levels, enhanced oxidative stress responses, and counterintuitively reduced virulence by downregulating key virulence factors (58). In our report, the most pronounced phenotypic effect observed was the reduced infectivity of LmxTGT2-KO parasites. However, none of the known virulence factors were among the most affected proteins, preventing a straightforward explanation for the observed decrease in infectivity. Notably, a substantial proportion of the significantly affected genes encode hypothetical proteins or proteins of unknown function, which leaves open the possibility that they may play important roles in *Leishmania*-specific differentiation or infection. On the other hand, GO term analysis revealed processes associated with RNA metabolism, which may suggest a broader impact of Q-tRNA depletion on translation.

Interestingly, the *LmxTGT2*-KO parasites, caused significantly smaller ear lesions in infected mice as compared to those infected with WT or AB strains. However, this reduction in lesion size occurred despite comparable parasite burden, suggesting that Q-tRNA deficiency impairs the parasite’s ability to elicit an inflammatory response. This finding is consistent with our *in vitro* macrophage infection experiments, in which Q-tRNA-depleted parasites exhibited reduced survival and lower infectivity. The outcome of *Leishmania* infection is largely determined by the host–parasite interaction, so the severity of the disease can vary. Therefore, the larger lesions observed in control infections may result from an exacerbated immune response rather than a higher parasite load (59–61). Indeed, the various strains in this study showed comparable levels of IgG1, but IgG2a levels were significantly lower in LmxTGT2-KO-infected mice. As a result, the IgG2a/IgG1 ratio, a marker of pro-inflammatory, IFN-γ-mediated Th1 responses (48, 49), was significantly reduced, indicating a weakened immune activation in the absence of Q-tRNAs. This may explain the significantly smaller lesions observed in mice infected with the KO parasites.

In conclusion, our study provides the first direct evidence that queuosine tRNA modification regulates infectivity in the eukaryotic pathogen *L. mexicana* through codon-biased translational control. Q-tRNA levels increase significantly during differentiation from the insect into the mammalian-infective stage, and loss of Q modification via LmxTGT2 knockout impairs protein synthesis, leading to a codon-specific imbalance in the proteome that affects parasite virulence. These findings establish Q as an additional, novel layer of gene expression regulation in different life-cycle stages of *Leishmania*, providing new insight into how these parasites fine-tune translation to adapt to host environments in the absence of classical transcriptional control.

## DATA AVAILABILITY

The data underlying this article are available in the article and in its online supplementary material.

## SUPPLEMENTARY DATA

Supplementary Data are available online.

## ACKNOWLEDGMENTS

This work was supported by the Czech Science Foundation grants 20-11585S, (to Z.P.), Lead Agency grant 23-08669L (to Z.P.) and by ERDF/ESF and MEYS CR projects Centre for Research of Pathogenicity and Virulence of Parasites [No. CZ.02.1.01/0.0/0.0/16_019/0000759 to Z.P.] and RNA for therapy [CZ.02.01.01/00/22_008/0004575 to Z.P.], We acknowledge CIISB, Instruct-CZ Centre of Instruct-ERIC EU consortium, funded by MEYS CR infrastructure project LM2023042, at the CEITEC Proteomics Core Facility. Computational resources were provided by the e-INFRA CZ project (ID:90254), supported by MEYS CR. We thank Prof. Yurchenko (University of Ostrava) for the pLEXSY constructs and *L. mexicana* Cas9 cell line, Anusha Kote Raveendra for the graphical abstract, and Prof. Alfonzo (Brown University) for his insightful feedback on the manuscript.

## Author contributions

B.K., J.K., M.B., S.K., and Z.P. designed the research; B.K., J.K., M.B., S.K., T.P.F., and Z.P. performed the research; B.K., J.K., M.B., S.K., T.P.F., A.S., and Z.P. analysed the data; and B.K., J.K. and Z.P. wrote the paper.

