## SUPPLEMENTARY MATERIAL 1 for "Codon biased translation mediated by Queuosine tRNA modification is essential for the virulence of *Leishmania mexicana*"

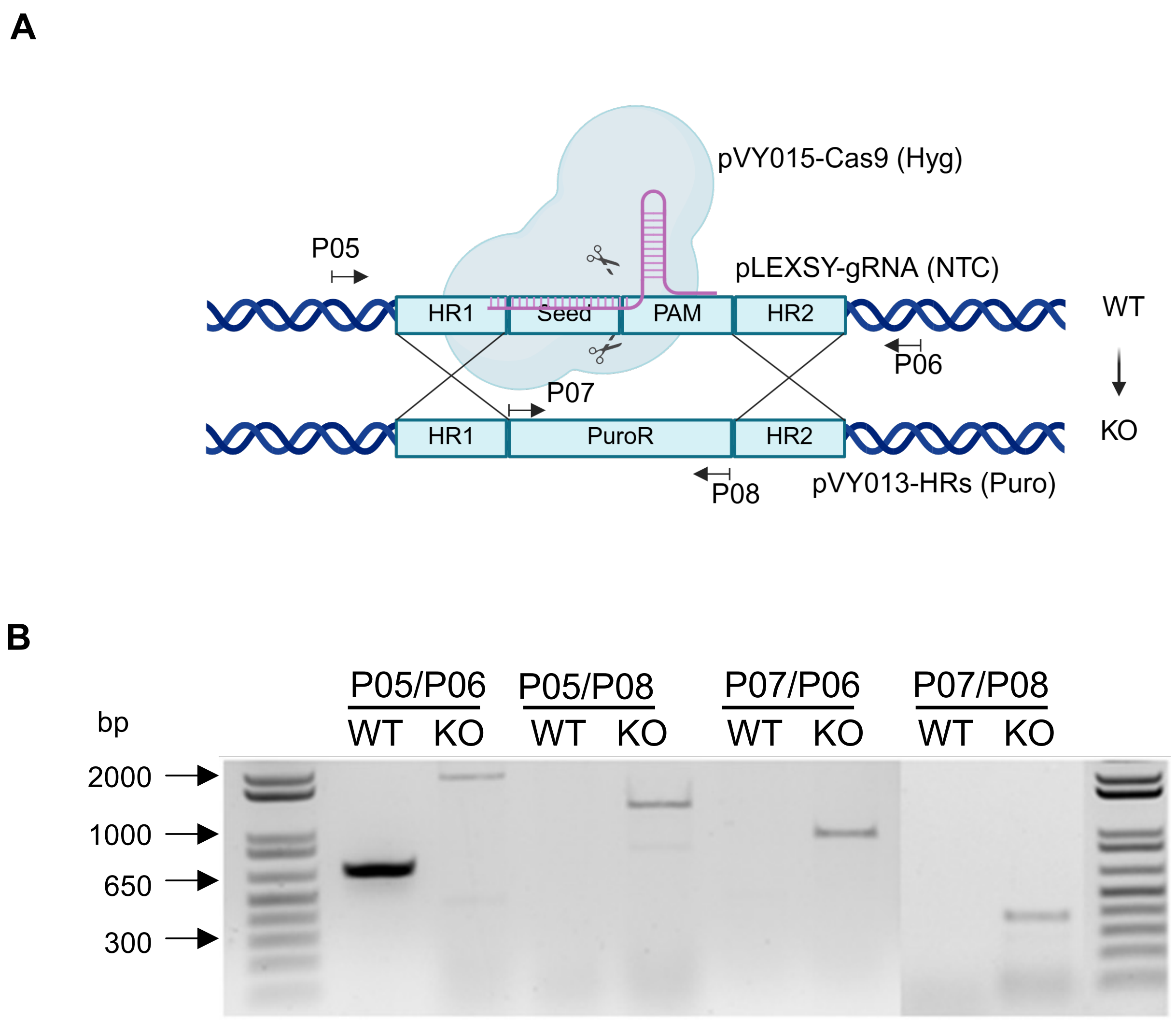


**Suppl. Fig. 1**

**Supplementary Figure 1. Design and confirmation of LmxTGT2 knockout in *L. mexicana***

**(A)** Schematic illustration of the strategy used to generate and validate LmxTGT2 knockout (KO) cells. Positions of PCR primers used for genotyping are indicated. **(B)** PCR-based amplification followed by DNA electrophoresis reveals the presence or absence of specific genomic sequences in wild-type (WT) and knockout (KO) cells. Product sizes (in base pairs) are indicated to confirm the specificity of the knockout strategy.

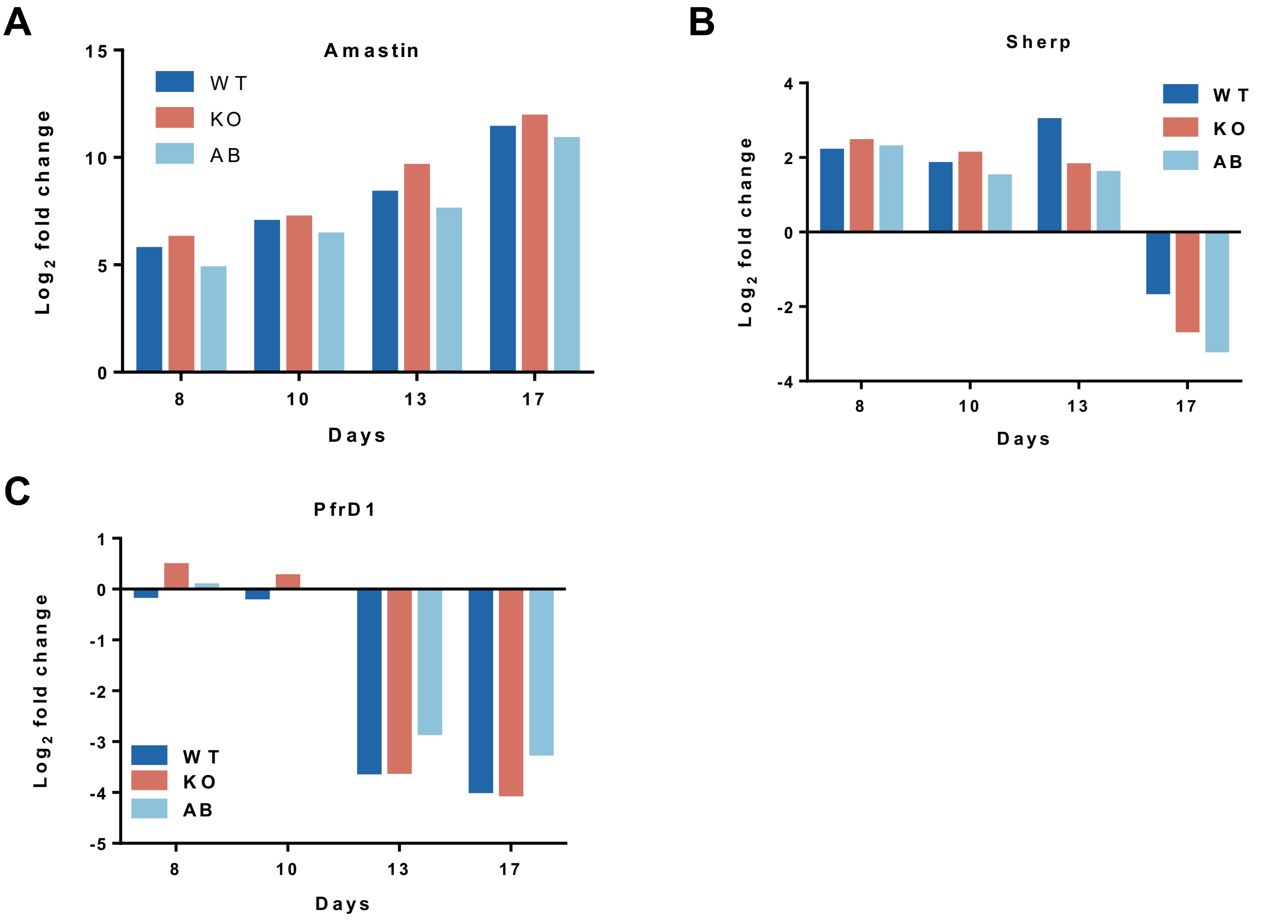


**Suppl. Fig. 2**

**Supplementary Figure 2. Proteomic analysis of stage-specific marker expression during differentiation**

**(A–C)** Proteomic expression levels of stage-specific markers Amastin, PfrD1, and Sherp during *L. mexicana* differentiation.

**A**

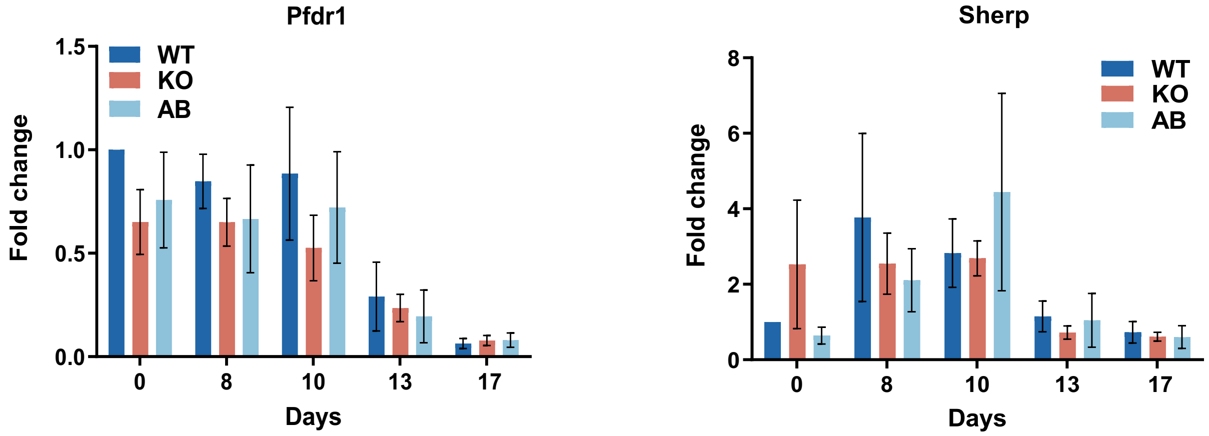


**Suppl. Fig. 3**

**Supplementary Figure 3. Depletion of Q-tRNAs does not affect stage-specific marker expression**

**(A)** qPCR analysis of PfrD1 and Sherp, markers of the promastigote and metacyclic stages, during *L. mexicana* differentiation.

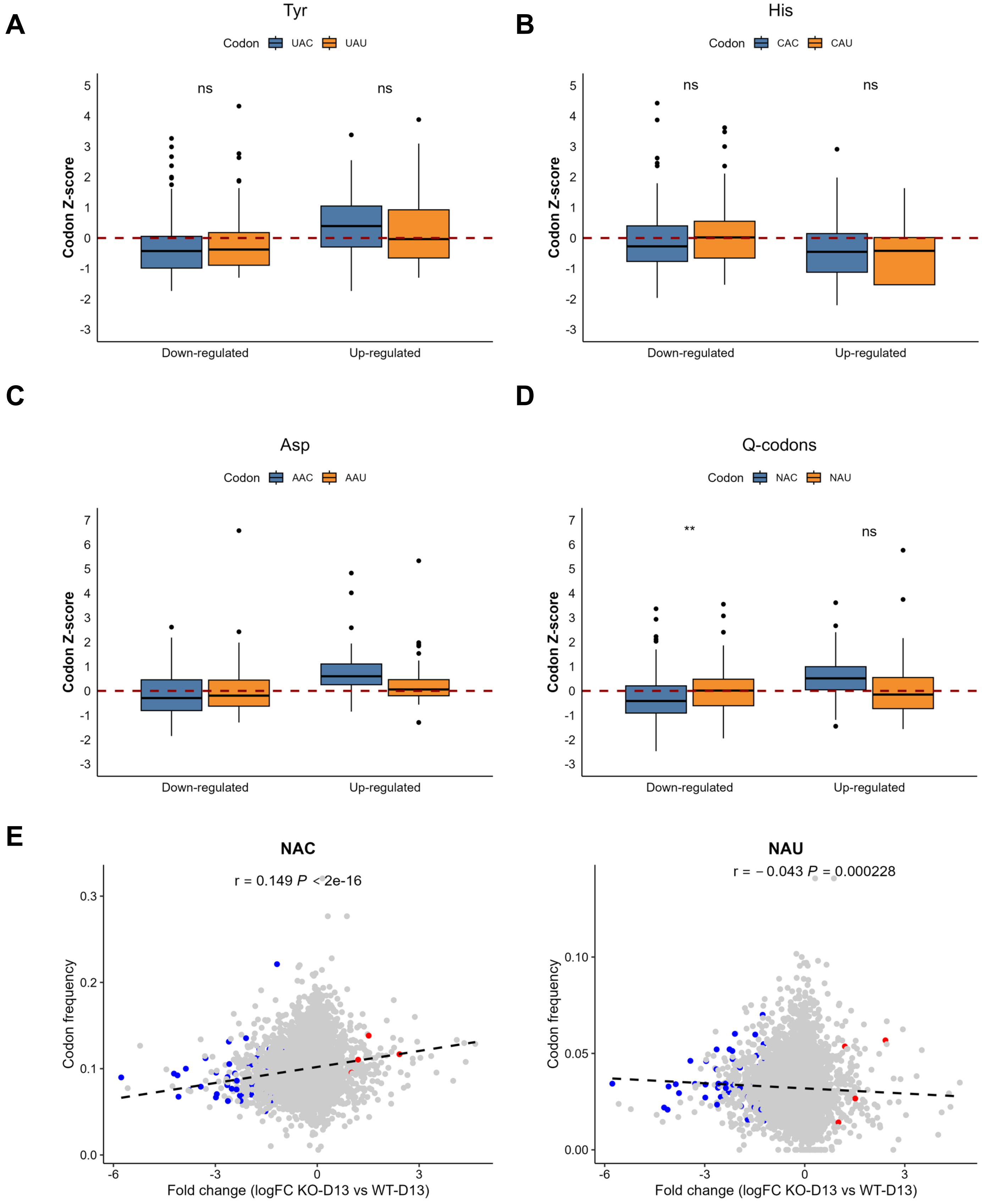
A
Tyr
Codon UAC UAU
ns
ns
Codon Z-score
Down-regulated
Up-regulated
B
His
Codon CAC CAU
ns
ns
Codon Z-score
Down-regulated
Up-regulated
C
Asp
Codon AAC AAU
Codon Z-score
Down-regulated
Up-regulated
D
Q-codons
Codon NAC NAU
**
ns
Codon Z-score
Down-regulated
Up-regulated
E
NAC
r = 0.149 P < 2e-16
Codon frequency
Fold change (logFC KO-D13 vs WT-D13)
NAU
r = −0.043 P = 0.000228
Codon frequency
Fold change (logFC KO-D13 vs WT-D13)

**Suppl. Fig. 4**

**Supplementary Figure 4. Regulation of translation via codon bias in the absence of LmxTGT2 (day 8)**

**(A–C)** Codon frequency of Asn, Tyr, and His, shown as Z-scores, in upregulated and downregulated proteins in LmxTGT2-KO cells at day 8, relative to wild-type (WT) and add-back (AB). **(D)** Codon frequency of all NAC codons, shown as Z-scores, in upregulated and downregulated proteins, compared to NAU codons at day 8. The centre line indicates the median, the box shows the interquartile range (upper and lower quartiles), and individual dots represent outliers (minimum and maximum values). **(E)** Pearson correlation between NAC and NAU codon frequency and protein abundance in LmxTGT2-KO cells at day 8, relative to WT and AB. The Pearson correlation coefficient (r) with *P* value and linear regression line (black) are indicated. The data are based on three independent biological replicates used for proteomic analysis and were analysed using Student's *t*-test. Significant differences are indicated as follows: (**$P < 0.01$, * $P < 0.05$, ns = not significant).

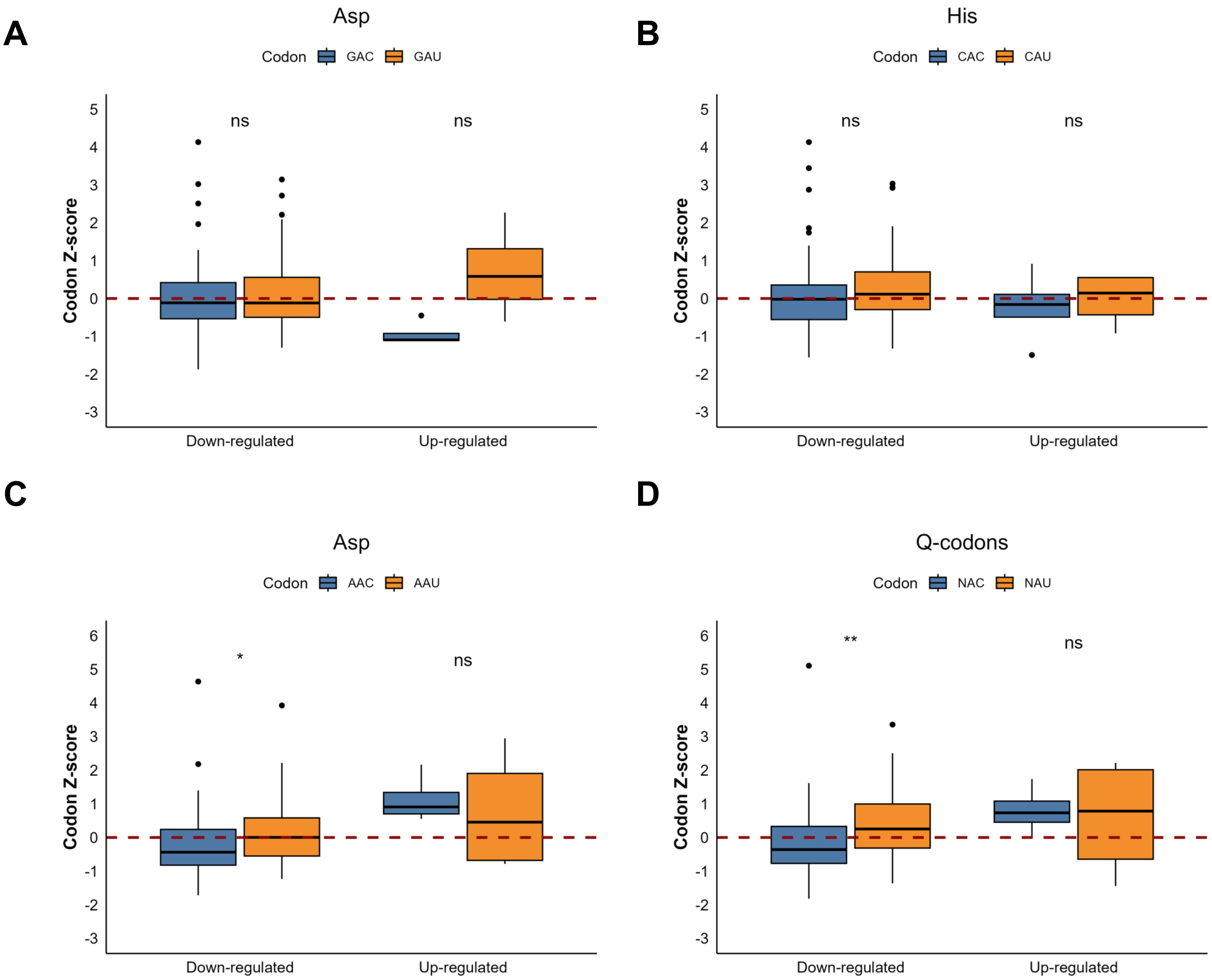
A
Asp
Codon GAC GAU
ns
ns
Codon Z-score
Down-regulated
Up-regulated
B
His
Codon CAC CAU
ns
ns
Codon Z-score
Down-regulated
Up-regulated
C
Asp
Codon AAC AAU
*
ns
Codon Z-score
Down-regulated
Up-regulated
D
Q-codons
Codon NAC NAU
**
ns
Codon Z-score
Down-regulated
Up-regulated

**Suppl. Fig. 5**

**Supplementary Figure 5. Regulation of translation via codon bias in the absence of LmxTGT2 (day 13).**

**(A–C)** Codon frequency of Asn, Asp, and His codons, shown as Z-scores, in upregulated and downregulated proteins in LmxTGT2-KO cells at day 13, relative to wild-type (WT) and add-back (AB). **(D)** Codon frequency of all NAC and NAU codons, shown as Z-scores, in upregulated and downregulated proteins in LmxTGT2-KO cells at day 13, relative to wild-type (WT) and add-back (AB). The center line indicates the median, the box shows the interquartile range (upper and lower quartiles), and individual dots represent outliers (minimum and maximum values). The data are based on three independent biological replicates used for proteomic analysis and were analysed using Student's t-test. Significant differences are indicated as follows: (***P* < 0.01, * *P* < 0.05, ns = not significant).

| Oligonucleotide | sequence (5'-3') | Description |
|---|---|---|
| **P01** | GAAGATGGTGCAGGTAAGCGgttttagagctagaaatagcaagttaaaa | LmxTGT2-KO generation (seed, sgRNA) |
| **P02** | CGCTTACCTGCACCATCTTCtccatcactgagcatacggtcaagacaagg | LmxTGT2-KO generation (seed, U6 promoter) |
| **P03** | GTGCGACTGCTTCACCTGCAAGCGGCACTaggcttgcacccaggctc | LmxTGT2-KO generation (5' TGT2 HRs) |
| **P04** | GAATGTCGCTGTTCATTTCCTGAACGGTcctccctatctctctctccgctc | LmxTGT2-KO generation (3' TGT2 HRs) |
| **P05** | ACCACGGTAGCCGGTGATAA | LmxTGT2-KO verfification (LmexTGT2 F) |
| **P06** | ACCACCTGCGCAAGATTATG | LmxTGT2-KO verfification (LmexTGT2 R) |
| **P07** | ATGACCGAGTACAAGCCCAC | LmxTGT2-KO verfification (Puromycine F) |
| **P08** | GAGGCCTTCCATCTGTTGCT | LmxTGT2-KO verfification (Puromycine R) |
| **P09** | GTGGTCCTTCCGGCCGGAATCGAA | Northern blot: tRNA-Tyr |
| **P10** | GTGGTCCTTCCGGCCGGAATCGAA | Northern blot: tRNA-Asp |
| **P11** | CTCCTCCCGTTGGATTCG | Northern blot: tRNA-Asn |
| **P11** | CGGACCCGGGTATTCAGAG | Northern blot: tRNA-His |
| **P12** | GTTCCGGAAGTTTCGCATAC | Northern blot: SL RNA |
| **P13** | GTTGTACACCGTCAGCTCCA | RT PCR: PfrD1 F |
| **P14** | CAAGGAGAACGAGGAGATGC | RT PCR: PfrD1 R |
| **P15** | CAAGGAGAACGAGGAGATGC | RT PCR: Sherp F |
| **P16** | GTTGTACACCGTCAGCTCCA | RT PCR: Sherp R |
| **P17** | TTCCTCGCGTTTCTCTTTGT | RT PCR: Amastin F |
| **P18** | AAAATGGAAATGACCGCAAG | RT PCR: Amastin R |
| **P19** | AACCAGTTCACGAAGGTGCT | RT PCR: 60S F |
| **P20** | CTTCGTCGACACCGTCTTCT | RT PCR: 60S R |

**Suppl. Table 1**

**Supplementary Table 1.**

List of oligonucleotides used for cell line generation, PCR validation and Northern blotting.
